# Experimental community ecology in decline: A call to embrace technology

**DOI:** 10.1101/2025.10.22.683879

**Authors:** Paulina A. Arancibia, Nerea Abrego, Peter Morin, Tomas Roslin, Otso Ovaskainen

## Abstract

Community dynamics are complex and thus challenging to infer from observational data alone. Experiments, with their ability to control variables and isolate mechanisms, are a powerful tool for uncovering the causal processes that drive community dynamics. They therefore allow us to move beyond correlations and to directly test theoretical predictions. Yet, because experiments are often logistically demanding and resource-intensive, they are less frequently employed than observational approaches in community ecology. Here, we trace the past three decades of experimental research in community ecology through a systematic literature review. We focus on the motivation behind experiments, their links to ecological theory, the types of questions they address, their scale, and the methods used to do this. Our results corroborate the historically tight relationship between experiments and ecological theory and document a gradual increase in experimental complexity —particularly related to the use of molecular methods. However, persistent gaps remain in the taxa and ecosystems studied, with aquatic ecosystems, fungi, and microbes still underrepresented compared to terrestrial plants and animals. Moreover, experiments are still limited in their spatial and temporal scale; they are typically short-term, local, and reliant on manual methods. The integration of high-throughput technologies with experimental workflows is still in its infancy, even though they are increasingly common in biomonitoring. To illustrate the potential of such tools in experimental research, we present a proof-of-concept study. It shows how automated technologies can be incorporated at different stages of the experimental workflow to expand the scale of experiments while reducing the reliance on human labor and potentially lowering financial costs. We conclude that many of the long-lasting biases and challenges in experimental community ecology could be addressed by combining technological innovations with broader collaboration among research groups. Coordinated networks, standardized protocols, and the integration of long-term and large-scale experimental designs can substantially improve in situ replication as well as cross-site comparability. Such efforts are essential for developing a more comprehensive mechanistic understanding of community dynamics across diverse ecosystems.

## 1. Introduction

### 1.1. Advancing community ecology through experiments

Community ecology aims to understand the patterns and processes emerging from multispecies systems. There are myriads of assembly processes that act and interact at different scales to create variation in community composition, starting from species interactions at the smallest scales to phylogeographic processes at the largest scales (Brown et al. 2004, Götzenberger et al. 2012, Vellend 2016, Keddy and Laughlin 2021). While community ecology aims to disentangle all these processes to gain a mechanistic understanding of community assembly and community dynamics, reaching this aim using observational data alone is difficult. Observational studies can provide valuable insights that help to formulate hypotheses about the potential drivers of the observed patterns (Underwood et al. 2000, Travis 2020, Byrnes and Dee 2025), especially for communities for which manipulative approaches are difficult to implement (Morin 2011). However, observations alone are not enough to infer a process from a pattern (Münkemüller et al. 2020). The main challenge is that multiple processes can lead to similar —or even identical— biodiversity patterns (Cale et al. 1989, Ovaskainen et al. 2019). Thus, the only robust way of gaining conclusive evidence about assembly processes, and ultimately achieving a predictive understanding of community dynamics, is through experimental research. To evaluate the progress of community ecology as a field, it is crucial to understand the advancements in experimental approaches within community ecology.

Experiments provide the opportunity to bridge theoretical and empirical research in community ecology. Since theory, and mathematical models in particular, can suggest the rules that drive different assembly processes, experiments are a powerful way to test theoretical predictions by evaluating the roles of each component both in isolation and combined. Experiments allow us to simplify scenarios by controlling for the abiotic and biotic environment in a way that is not possible in non-manipulative observational studies, targeting specific community-level processes (e.g., Tilman and Downing 1994, Kaunzinger and Morin 1998, Arancibia 2024, Abrego et al. 2025). At the same time, experiments can also feed theory by providing data about the starting conditions or the parameters to make mathematical models more realistic (e.g., Petermann et al. 2008, Carrara et al. 2015, Shen et al. 2023). However, the cost of and the often elaborate logistics behind community-level experiments can restrict their scope and/or scale of inquiry (Sasaki et al. 2025). This is especially true for species-rich communities where species interact with each other and with the environment in complex and unknown ways. Thus, despite the advantages experiments offer for robustly addressing topical questions in community ecology, experiments may often not be the first choice for empirical ecologists.

### 1.2. The growth of community ecology as a core field in ecology

As a relatively young area of modern biology, community ecology is undergoing a swift, transformative change. Twenty years ago, the field was described as “a mess with so much contingency that useful generalizations are hard to find” (Lawton 1999). Yet, these last two decades have brought rapid empirical and theoretical advances. Two ingredients represent important steps in the progress of community ecology: a series of large-scale comparative studies exposing generalities in patterns of biodiversity across ecosystems and taxa (Hillebrand and Blenckner 2002, Belmaker and Jetz 2012, Soininen 2014, Linquist et al. 2016, Nishizawa et al. 2022), and the development of unified theoretical frameworks (Chase and Leibold 2003, Leibold et al. 2004, Vellend 2010, Keddy and Laughlin 2021). As a result of this growth, hypotheses in, for example, community assembly have proliferated; consequently, the next logical step for advancing the field is to test these data-driven and theoretical hypotheses experimentally. Fifteen years ago Logue et al. (2011) concluded that experimental research in metacommunity ecology was highly biased towards those processes that are easier to test, especially environmental filtering, i.e., the effect of the abiotic environment on community structure. However, moving towards a predictive understanding of community ecology requires an integrative understanding of the roles of all processes, including biotic filtering, dispersal, stochastic, and evolutionary processes. Thus, experimental studies should be motivated by a broader range of processes than environmental filtering. Likewise, community ecology frameworks are often biased towards the best-known species groups, such as larger plants and animals (Götzenberger et al. 2012, Keddy and Laughlin 2021). This bias is partially due to the large knowledge gaps in microbial taxonomy and the specialized expertise required to study them, which has caused microbial community ecology to lag behind that of plants and animals (Nemergut et al. 2013). A general understanding of community dynamics requires the inclusion of all organismal groups, including microbes.

### 1.3. A brief history of experimental approaches in community ecology

Although community ecology initially emerged as a largely observational and descriptive endeavor, experiments gradually became important as a means of testing hypotheses about causal processes. As in other branches of ecology, experimental approaches in community ecology range from controlled laboratory studies to manipulative field experiments. These manipulations are typically categorized as either press or pulse disturbances (Bender et al. 1984). Press disturbances involve sustained, continuous alteration of environmental conditions over time, whereas pulse disturbances are short-term, discrete events that temporarily modify the environment before conditions return to their baseline.

Laboratory experiments in community ecology, while limited in ecological realism due to their simplified species compositions and controlled settings, offer significant advantages. The main advantage is that they allow for controlling abiotic variables (e.g., light, water, nutrients) that in real ecosystems may vary unpredictably across space and time. Laboratory microcosm experiments (i.e., small experimental ecosystems) have boosted the formulation and empirical testing of foundational principles in community ecology. Microcosm experiments typically include only a few species of small organisms (e.g., algae, bacteria, arthropods) and small resource patches (e.g., multi-well plates, petri dishes, or flasks). For instance, Gause demonstrated the ‘competitive exclusion principle’ using experiments with two *Paramecium* species that had identical resource use (Gause 1934). In another classic microcosm experiment, Huffaker (1958) designed a system with a predatory mite and a prey mite that fed on oranges (used as resource patches). By manipulating the spatial arrangement of these patches, he was able to produce predator-prey cycles analogous with the dynamics predicted by Lotka-Volterra models. Later, Tilman (1977) cultured two species of diatoms under a gradient of two resources to investigate the role of resource competition in shaping community structure, while evaluating the ability of mechanistic models to predict competitive outcomes. This pivotal experiment helped to develop the resource-ratio hypothesis, which posits that species coexistence is mediated by trade-offs in resource acquisition under varying nutrient conditions.

Compared to laboratory experiments, field experiments offer a higher degree of ecological realism, albeit at the cost of reduced control over biotic and abiotic variables. Conducted in natural ecosystems, these studies capture more natural and complex variation than laboratory studies, and allow for testing hypotheses on real communities consisting of many species — without being constrained by the size or number of the organisms considered. Field experiments can be conducted ex-situ, for example translocating habitat patches outside their original environment, or in situ by manipulating the habitats or the species composition within their original environment.

Field experiments have also greatly contributed to the formulation and testing of foundational principles in community ecology. They started to gain popularity in the 1960s as the main means to test hypotheses regarding the importance of biotic interactions in structuring communities (Bertness et al. 2019). Using the rocky intertidal zone as their stage, Joseph Connell, Robert Paine, and their students initiated a wave of hypothesis-driven experimental studies that shaped many foundational concepts in community ecology. Through field experimental manipulations, Connell helped shape our understanding of competition and niche partitioning (Connell 1961a, 1961b). Likewise, the work of Robert Paine set the basis for the understanding of the role of top-down control on community structure (Paine 1966), and coined the “keystone species” concept (Paine 1969), among others. Experiments by Paul Dayton in the rocky intertidal and by Wayne Sousa in intertidal boulder fields demonstrated the influence of interactions, disturbance, and priority effects on successional dynamics (Dayton 1971, Sousa 1979). In another landmark field experiment in community ecology, Simberloff and Wilson (1969) monitored species recolonization patterns in mangrove islands after artificial defaunation. This experiment provided the key early empirical test of the predictions of the Theory of Island Biogeography (Macarthur and Wilson 1967), and showed that islands can reach a new equilibrium with a species richness similar to the pre-defaunation state, supporting the theory’s core concepts. Notably, most of these foundational experiments were performed in marine ecosystems, a trend that seems to have shifted in recent decades.

While the contributions of experiments to the development of community ecology are indisputable, especially in its early era, concerns have been raised about certain limitations in contemporary experimental approaches. It has been argued that experimental research often falls short of manipulating the relevant biotic and abiotic drivers of community assembly at realistic levels of variation, over relevant ecological scales of space and time, and of integrating the responses of large numbers of species simultaneously. Such shortfalls may then confound the interpretation of direct and indirect effects (Keddy and Laughlin 2021).

### 1.4. New technologies for community ecology research

Technological advances have changed the way ecologists collect data. In the last decade alone, the use of technology, particularly for monitoring changes in biodiversity, has virtually exploded (Pawlowski et al. 2021, Besson et al. 2022, Hartig et al. 2024). The use of high-throughput methods, such as audio recording, camera traps, and environmental DNA (eDNA), has become increasingly popular. In particular, the use of eDNA has revolutionized our ability to survey previously poorly known and cryptic species groups such as bacteria (Garlapati et al. 2021), fungi (Abrego et al. 2024), and invertebrates (van Klink et al. 2022). As a result, observational studies in community ecology increasingly include data on these species-rich communities (Hartig et al. 2024). Yet, as highlighted in a recent review by Hartig et al. (2024), the use of data from high-throughput methods is still in its infancy, despite its many opportunities for advancing the understanding of fundamental community ecology. Automated methods are already facilitating the collection of large amounts of biodiversity data and reducing the reliance on traditional in-person observations. Building further on these gains, the use of other technologies such as robotics could extend the scale of experimental research beyond the practical limits of human involvement (Besson et al. 2022).

## 2. Review I: Has experimental community ecology evolved along with the growth of community ecology?

### 2.1. Methods for Review I

To evaluate whether the ability and approaches used to experimentally understand the complexity of community dynamics have changed over time, we explored the changes in the frequency and complexity of experimental approaches over time in community ecology through a literature review (see Figure 1). We used the terms “experiment” * “community ecology” in the ISI Web of Knowledge database, narrowing the results to studies published 1991-2023. The search yielded 8595 papers, which we ranked by relevance using the website’s built-in sorting tool. We used a stratified random sampling approach and selected the 10 most “relevant” studies per year. We analyzed 330 manipulative studies, and asked whether the ecological context, methodologies, or the focal points of experimental studies in community ecology have shifted in response to advances in biomonitoring technologies, as well as in empirical and theoretical understanding of community assembly. In particular, we evaluated shifts in 1) motivation, by assessing yearly changes in the proportion of experiments motivated by theory, patterns in observational data, or a knowledge gap in the specific study system; 2) question types, by evaluating changes in the community attribute targeted; 3) experimental design, by comparing shifts over time in the methods employed and the type of study system targeted; and 4) scale and complexity, by assessing whether the spatio-temporal scale and size of the community studied have increased over time (see Table 1 for the working definitions of each category). *A priori*, we expected a) a temporal shift towards studies that increasingly build on theoretical predictions and hypotheses informed by previous experimental research; b) a trend toward more experiments involving a larger set of community attributes, as driven by the integration of other subfields such as ecosystem ecology (Loreau 2010) and evolutionary ecology (Johnson and Stinchcombe 2007) with community ecology; c) a broadening of the experimental focus toward studies involving previously underrepresented taxa such as microbes, facilitated by technological advances used in biomonitoring; d) an increasing emphasis on more complex experiments (i.e. involving more variables) enabled by the development of new technology; e) a shift toward studies conducted at larger spatial scales and encompassing larger communities, as facilitated by technological advances which reduce the need for intensive human labor. To our surprise, only some of these trends were supported by our analysis.

**Figure 1.**
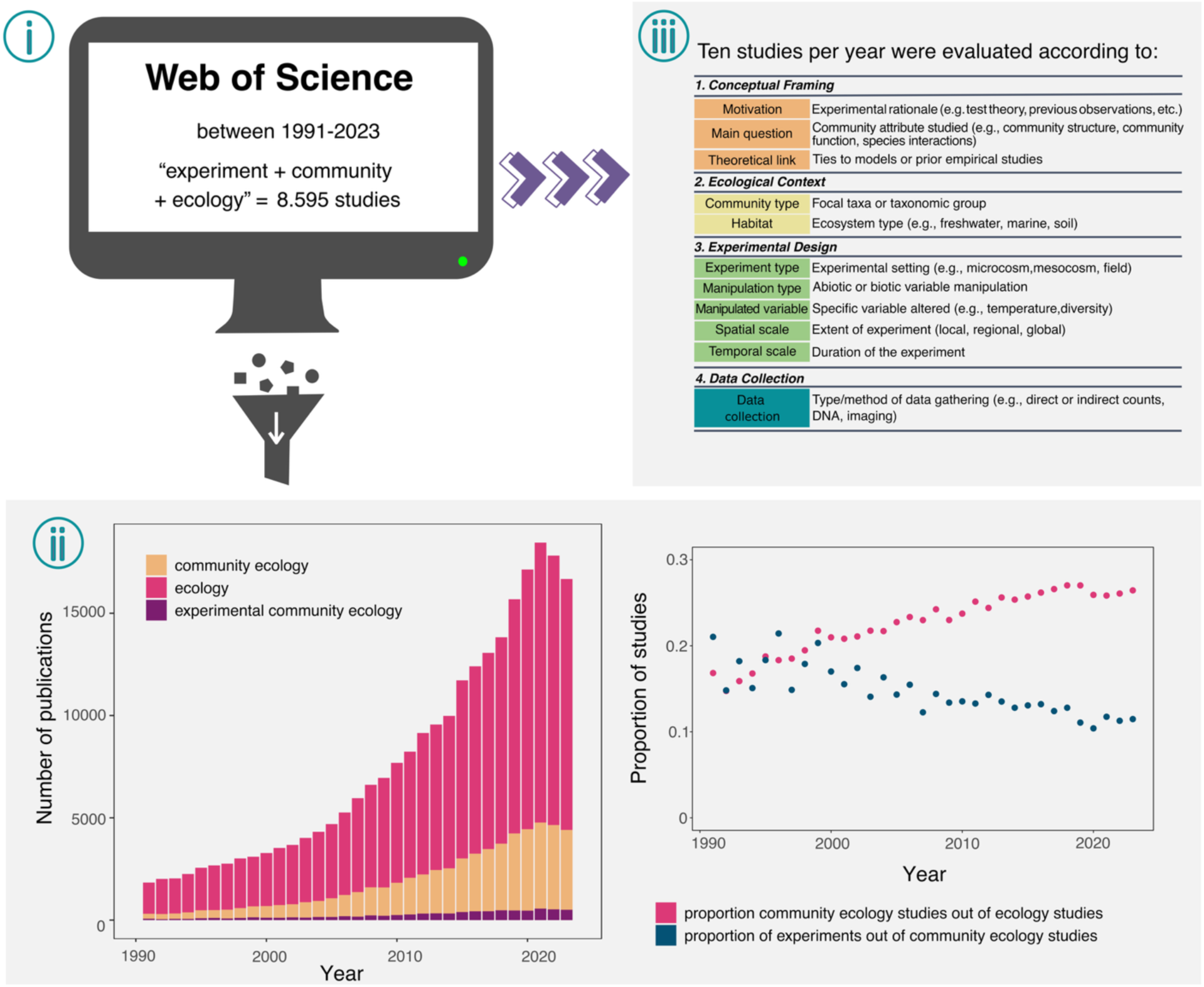
(i) Systematic review of experiments in community ecology using the terms “experiment” * “community ecology” in the ISI Web of Knowledge database. This search returned 8595 papers, which we ranked by relevance using the website’s built-in sorting tool. Using a stratified random sampling approach, we identified the 10 most relevant studies per year, with the primary selection criterion being that the study involved an explicit experimental manipulation. This sampling scheme ensured a balanced review of 330 studies published between 1991 and 2023. (ii) From each study, we tabulated information regarding 1) the conceptual framing of the experiment, their rationale, and the theoretical background of the study; 2) the ecological context of the study; 3) the experimental design implemented; and 4) the method used in data collection. (iii) The review revealed that while publications using the term “ecology”, and more specifically “community ecology” have steadily increased over the past three decades, the proportion of publications focusing on “experimental community ecology” has declined since the year 2000. Figure created by Paulina Arancibia.

**Table 1:**
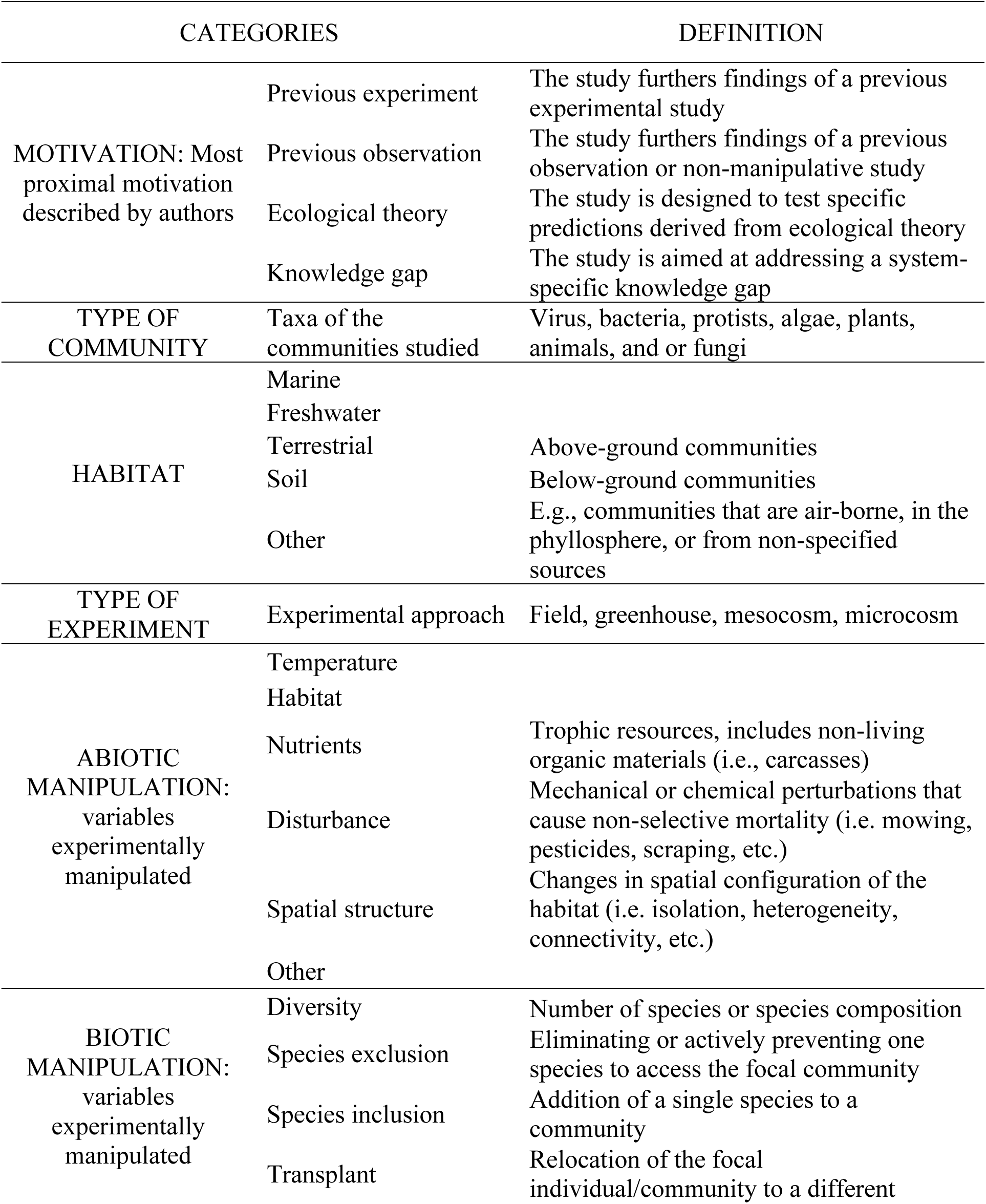
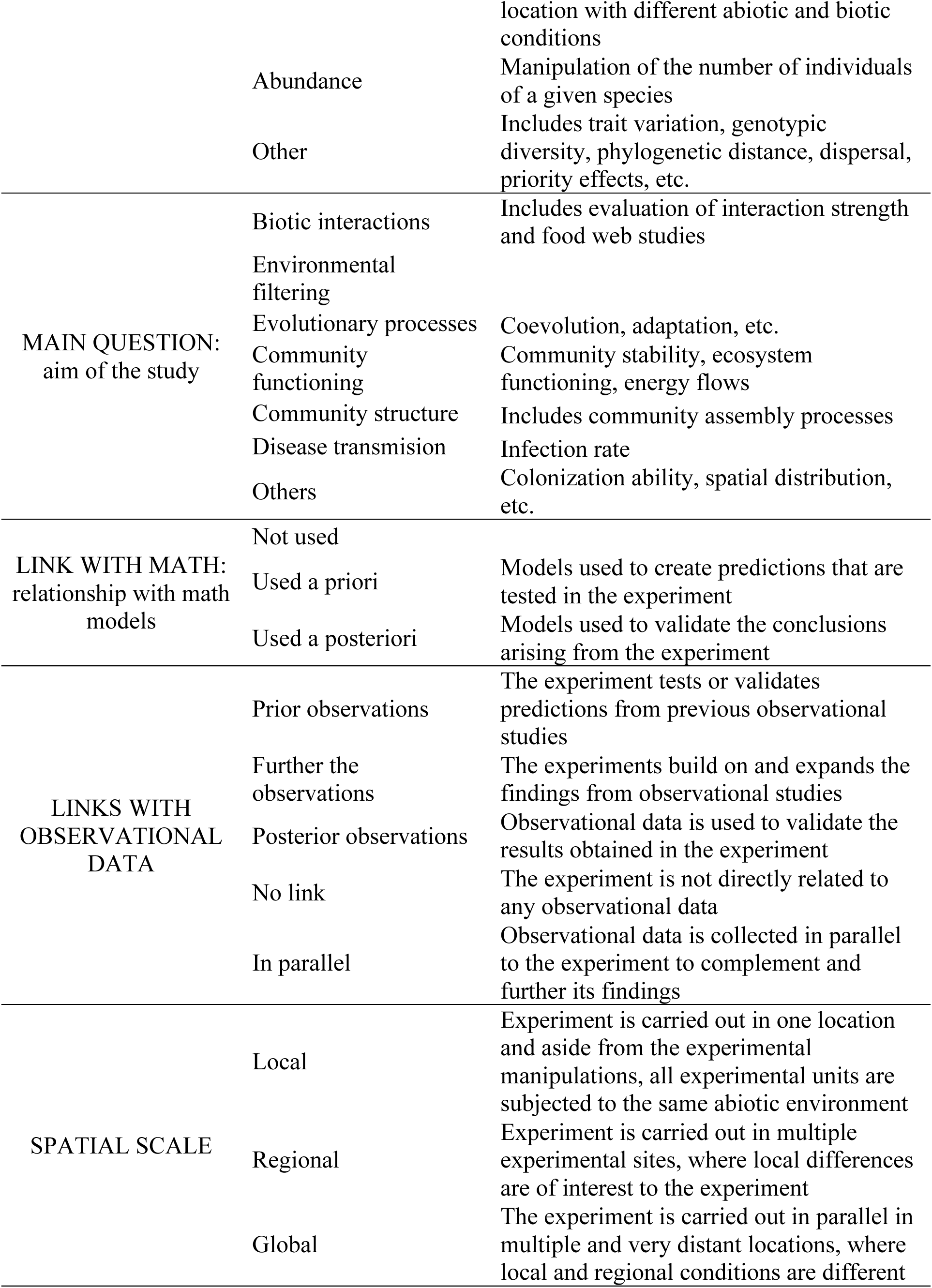
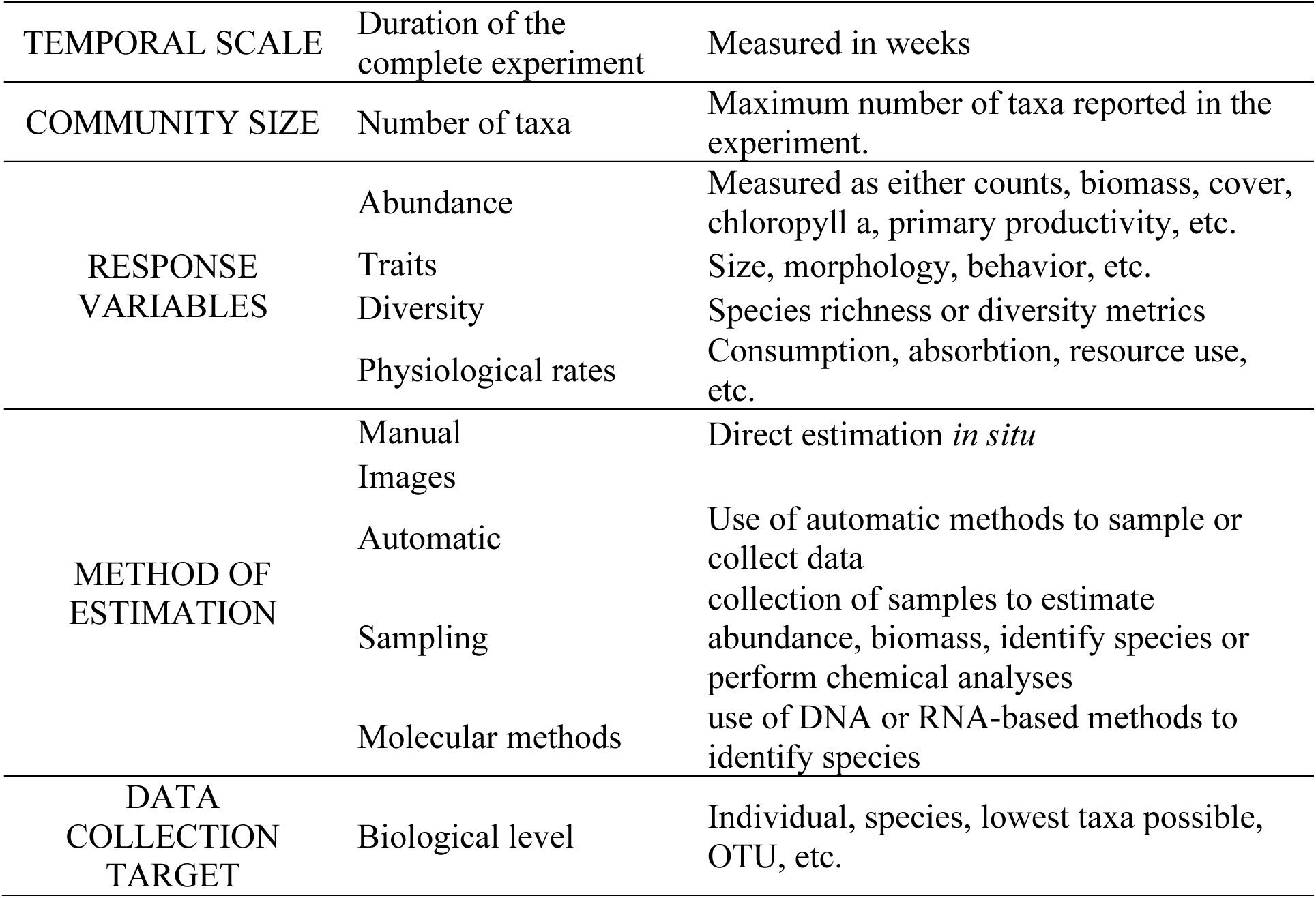
Definitions of the categories used to tabulate the information of the experimental studies included in the review.

### 2.2. Results of review I

#### Conceptual framing

Overall, directly testing ecological theory was the main motivation of experiments, followed by addressing system-specific knowledge gaps (Figure 2A). Across years, an average of 44% of the studies reviewed were motivated by testing or validating ecological theory, followed by filling a knowledge gap (38%), and testing hypotheses derived from previous experiments (11%) or observations (5%). While this pattern has remained stable over the years, the last decade has seen an increase in experiments motivated by hypotheses derived from previous studies and observations (Figure 2A).

**Figure 2.**
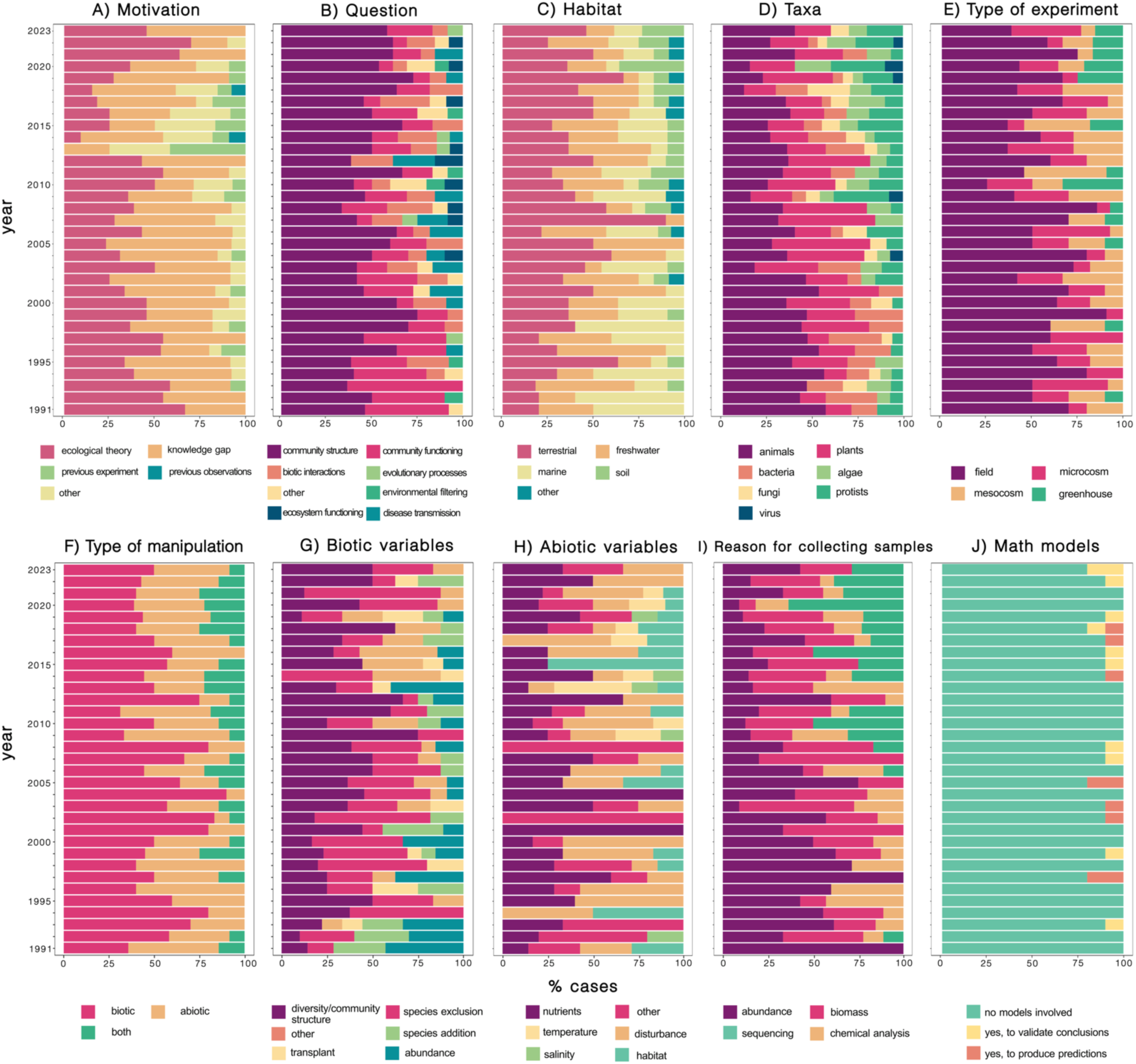
Results of the systematic review (Figure 1) of experiments in community ecology over the past three decades. *Motivation* (A) describes whether the primary purpose of the experiment was to test theory, clarify patterns observed in previous observational or experimental data, or fill a knowledge gap in the specific study system. *Question* (B) refers to the targeted response variable, e.g., whether the experiment assessed the influence of a treatment on community structure or ecosystem functioning. *Habitat* (C) describes whether the experiment was conducted in a terrestrial, marine, or freshwater context. *Taxa* (D) describes the targeted taxa at the level of kingdom. *Type of experiment* (E) describes whether the experiment was conducted in the field, microcosms, mesocosm, or greenhouse. *Type of manipulation* (F) describes whether biotic or abiotic variables were manipulated in the treatment. The specific variables that were manipulated are described under *Biotic variables (G)* and/or *Abiotic variables (H)*. For experiments that collected samples rather than measuring variables in situ, *Reason for collecting samples* (I) describes the purpose of collection. *Math models* (J) describe if and how the experiment was related to mathematical models. To account for studies reporting multiple responses, we show the distribution of studies among categories as percentages of cases.

Although experiments are often closely linked to ecological theory, the explicit use of theoretical/mathematical tools in experiments is infrequent —only in rare cases are mathematical models specifically used to validate and expand experimental findings (Figure 2J).

Changes in community structure are by far the most frequent community attribute studied via experiments, accounting for 52% of the studies across the years (Figure 2B). This is followed by experiments focusing on various aspects of community functioning, which account for 11% of the studies. However, after the 2000s, there has been a diversification of the focal community attribute studied. Experiments focusing on biotic interactions, disease transmission, ecosystem functioning, and evolutionary processes have increased over time, but still represent a small fraction of all studies combined (Figure 2B).

#### Ecological context

Over time, the focal ecosystem of experiments in community ecology has shifted. In the 90s, experiments in marine systems were generally more common than experiments conducted in terrestrial and freshwater systems, but over the last two decades, emphasis has increasingly turned toward terrestrial and freshwater systems (Figure 2C).

In terms of focal taxonomic groups, there is a persistent taxonomic bias in community ecology experiments, with larger, more easily enumerated animals and plants consistently being the most frequently studied groups (Figure 2D). However, in the last two decades, there has been a noticeable expansion in taxonomic coverage, with an increased representation of bacteria, protists and fungi. Similarly, experiments involving multiple taxonomic groups have become more frequent over time.

#### Experimental design

Field experiments are consistently the most prevalent type of experiment in community ecology, accounting for 57% of the studies across the years (Figure 2E). Microcosms and mesocosms account respectively for 19% and 16% of the studies across the years. After the 2000s, greenhouse experiments have also been included in community ecology, but still represent a minority (Figure 2E).

While biotic manipulations remain slightly more frequent than abiotic ones, there has been a growing trend toward experiments that simultaneously manipulate both biotic and abiotic factors (Figure 2F). Among abiotic variables, nutrient availability and disturbance regimes are the most frequently manipulated (Figure 2H), whereas biotic interventions most frequently involve changes in diversity or community structure, in particular species-exclusion experiments (Figure 2G). Moreover, most experiments rely on press perturbations, while only a small minority use pulse perturbations.

Experiments in community ecology are predominantly conducted at local spatial scales (Figure 3A). While regional-scale experiments are becoming more common over time, global-scale experiments are still rare. In terms of temporal scale, field experiments are by far the longest in duration, averaging 104 weeks. In contrast, studies in all other categories typically last around 20 weeks (Figure 3B). Importantly, though absolute measures of time —such as weeks or years— are only meaningful when considered in relation to the generational time of the focal organisms, and the ecological time scale of the processes under study.

**Figure 3.**
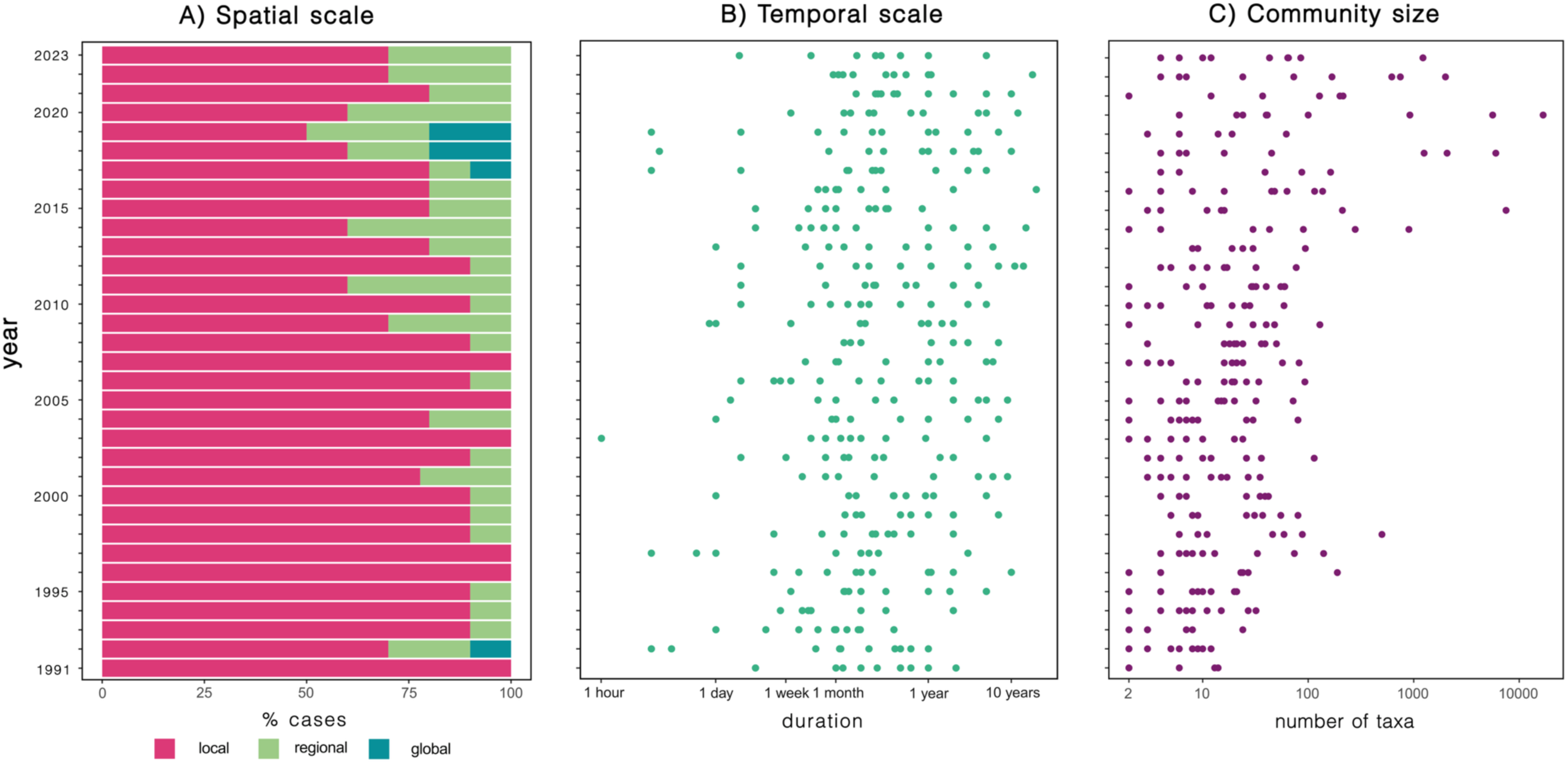
Evolution of “experimental complexity” in community ecology studies over the past three decades. *Spatial scale* (A) shows the proportion of experiments that we classified as local, regional or global. *Temporal scale* (B) shows the duration of the experiment, and *Community size* (C) the number of taxa studied, in both panels the dots correspond to individual experiments.

The complexity of communities targeted in experimental research has increased over time. Over the past decade, we observed a marked increase in the number of taxa included in experiments (Figure 3C), and this trend closely follows the rise in the use of molecular methods (Figure 2I).

#### Data collection

The last two decades have experienced a noticeable increase in the number of experiments using high-throughput sequencing techniques to characterize communities. This is also reflected in the level of biological organization recorded. In the last two decades, traditional taxonomic units such as individuals or species have been joined by operational taxonomic units (OTUs).

## 3. Review II: Has experimental community ecology embraced technological developments?

### 3.1. Methods for Review II

To better understand the role of technology in experimental studies, we reviewed another subset of 10 studies per year, for the years 2013-2022, selected using the same criteria as above. When a single study contained multiple experiments meeting our inclusion criteria, each qualifying experiment was counted and analyzed as a separate case. To tabulate the information, we divided the workflow of experimental research into three main steps: (1) experimental implementation, (2) primary data acquisition, and (3) data processing and extraction. We examined the methods used in each study to determine the proportion of experiments that employed automated technology.

### 3.2. Results for Review II

Our in-depth review of experimental methods revealed that in the last decade, the adoption of technological advances in experimental community ecology remains limited, despite their proliferation in biomonitoring. All (100%) of the experiments reviewed rely solely on manual methods during the experimental setup and maintenance phase, 5-36% use automated methods to collect data, and only 34% use some level of automation to process the data (Figure 4A). Here, sequencing and bioinformatics are the most prevalent high-throughput method employed (Appendix S1: Figure S1). Manual methods used to count, measure, and identify species account for over 73% of the methods used in these experiments (Figure 4B).

**Figure 4.**
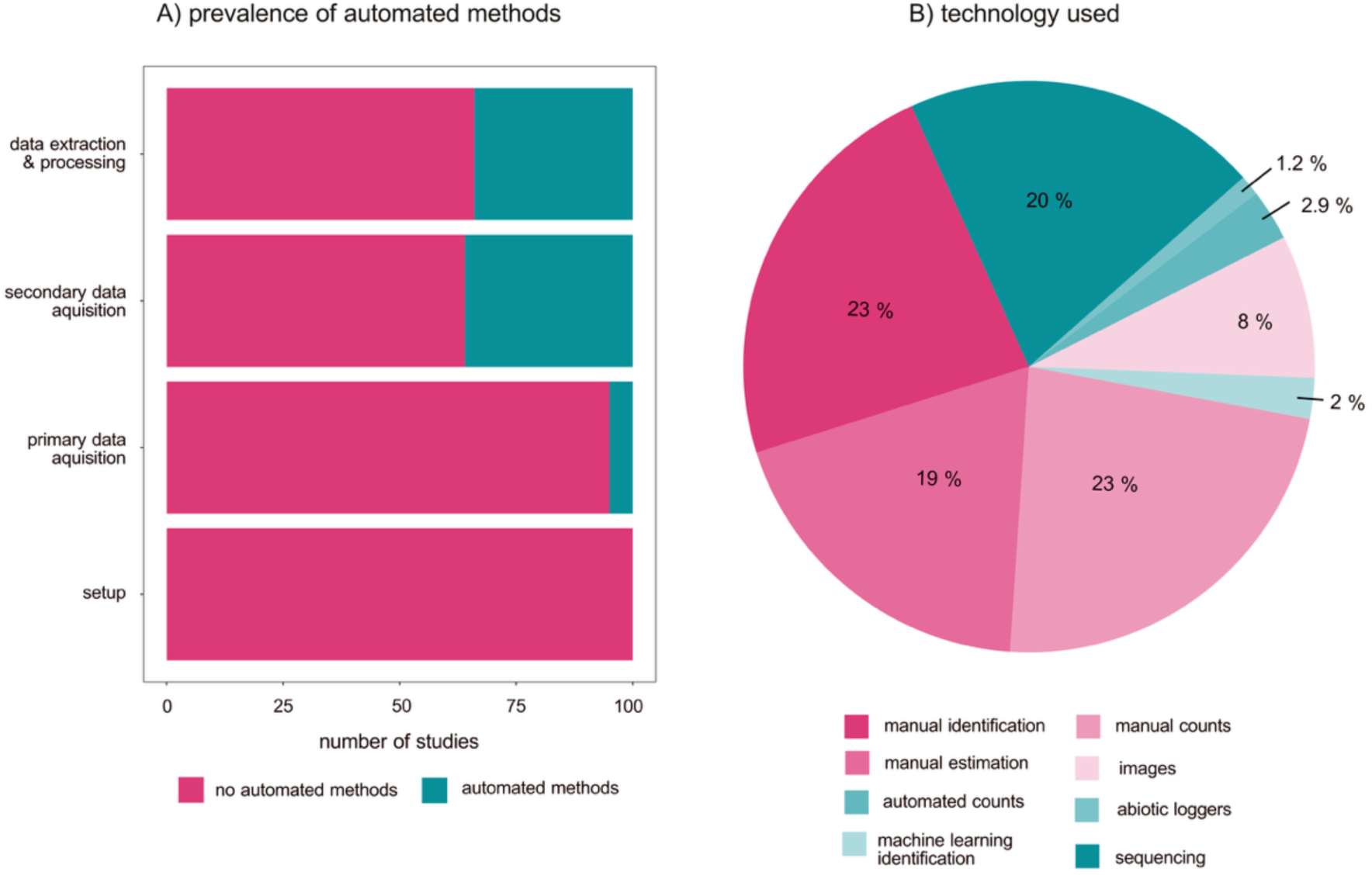
Results of an in-depth review of experimental methods in experimental community ecology: A) number of studies incorporating automated methods at different stages of the experimental workflow (shown as the number of studies in a given category, out of 100 studies included). B) Types of technological tools employed in experimental community ecology studies.

## 4. Proof of concept: Potential of automated methods to increase cost-efficiency in experiments

Motivated by the low use of automated methods revealed by the review, we attempted to understand the underlying reasons: is it because researchers are resistant to adopting new methodologies, or is it because the new technologies are not sufficiently reliable or cost-efficient? To address this question, we conducted a pilot study to compare the feasibility and efficacy of manual versus automated methods in microscopic counting of protists. This study system was selected as we had much prior experience with manual methods (Kaunzinger and Morin 1998, Arancibia and Morin 2022, Arancibia 2024) but no first-hand acquaintance with automated methods. Nonetheless, methods for counting protists from images (Pennekamp et al. 2015, 2017) and the availability of microscopes with robotic capabilities (Besson et al. 2022) suggested a high potential for automation. We compared the efficacy of manual versus automated methods as a function of the number of “counting units”, i.e., units for which the individuals are to be classified to their species and counted. For example, in the context of an experiment on the effects of spatial connectivity on metacommunities conducted by Arancibia (2024), one experimental unit consisted of a network of 24 connected communities. As the local communities were counted three times a week for one month, each experimental unit resulted in ca. 300 counting units, and having four replicates of two treatments resulted in a total of ca. 2500 counting units. Performing more complex experiments (as targeting several potential drivers with a higher level of replication) may then necessitate the inclusion of tens or hundreds of thousands of counting units.

The results of this pilot study (Figure 5) demonstrate the high potential of an automated method for large-scale experiments. First, after generating a small amount of calibration data by manual counting, it was possible to obtain reliable counts by the automated method. Second, while the initial cost of purchasing a microscope with robotic capabilities was substantial, automated methods spared the researcher’s time. With our pilot study, the break-point yielding identical financial cost between manual and automated methods was ca. 100,000 counting units. Beyond this point, the automated methods became orders of magnitude cheaper than the manual approach.

**Figure 5.**
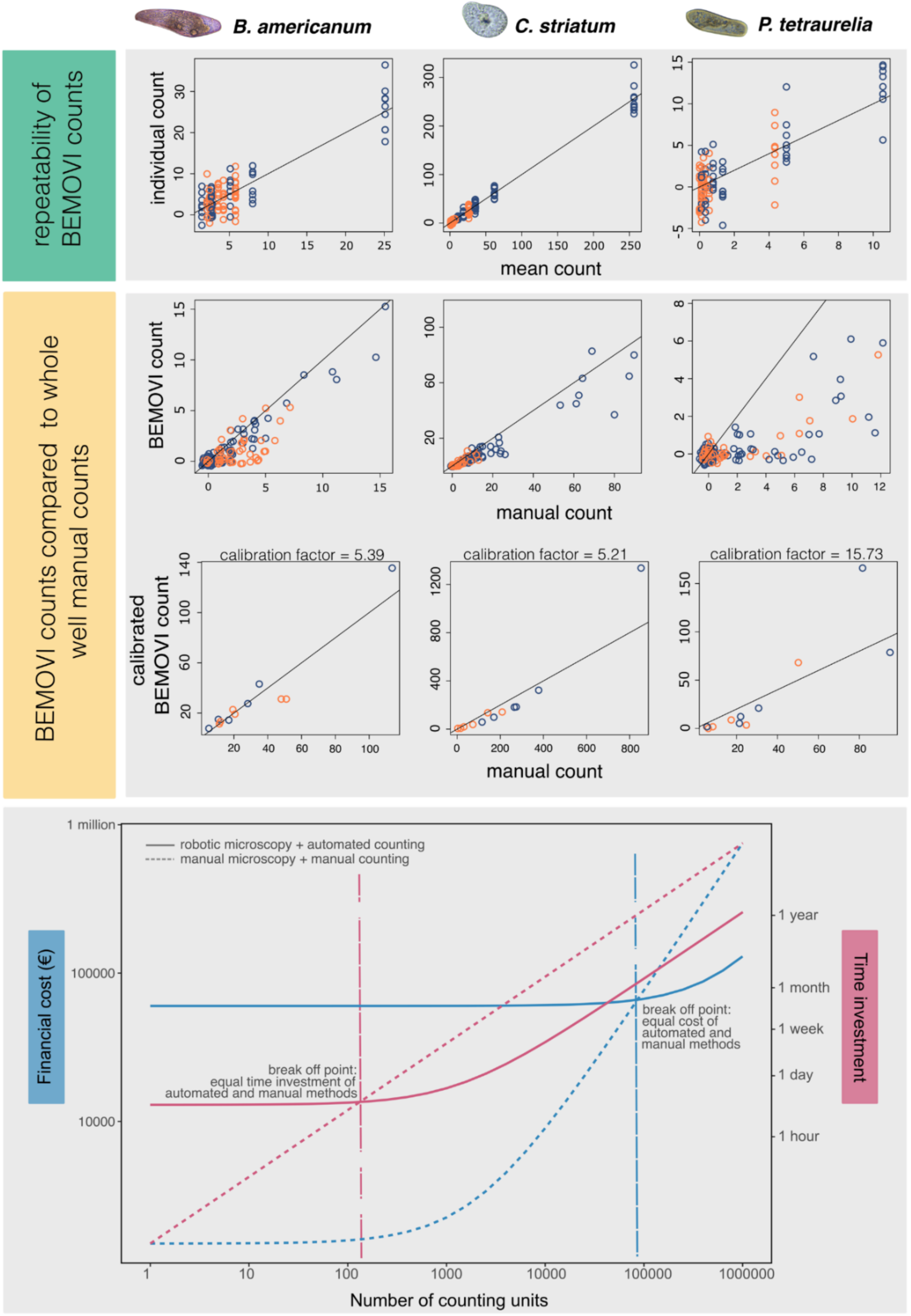
Top panels: results of a pilot study using aquatic protist microcosms to test the feasibility of a high-throughput experimental system that couples robotic microscopy with machine learning techniques to count and identify protists in single and mixed cultures. Green panels compare single video counts made using BEMOVI (Pennekamp et al. 2015) with the mean counts of the same sample (averaged over ten replicate counts). Yellow panels compare BEMOVI counts from videos with manual counts of the same video before (top) after (bottom) introducing a calibration factor that is derived from manual counting of 120 counting units. Blue circles represent single-species populations, and orange circles show multi-species communities. Lower panel: cost/time estimate shown up to a million counting units. The cost estimates include the price of a robotic microscope (€60,000) for the high-throughput approach and the price of a regular microscope (€1,500) for the manual approach. The cost of working time is budgeted according to the Finnish salary settings, and it refers to the actual cost for a project including side costs and overhead costs. The working time estimate is based on one hour of researcher time resulting in 40 manually counted units or 432 automatically counted units. Image credits: Paulina Arancibia.

## 5. Discussion

Our review shows a substantial increase in the interest in community ecology in recent decades, but an actual decrease in the proportion of studies involving experimental approaches. Since the ultimate goal of community ecology is to gain predictive insights into community assembly and dynamics, such a trend is highly unexpected. What causes it may be hard to establish. As contributing factors, we suggest the logistical complexity, extended timeframes, and higher risk of inconclusive or negative results associated with experiments than with observational studies. As observational studies are based on analyses of patterns, they can build from the accumulation of data on multiple ecosystems over time (Hampton et al. 2013, Zipkin et al. 2021). Perhaps this aspect of low risk may explain their growing prevalence in comparison to experimental approaches? The increasing reliance on observational data may also be spurred by their increased availability. Indeed, the rise of high-throughput approaches such as remote sensing, environmental DNA (eDNA), and other automated biodiversity monitoring tools has resulted in a virtual data deluge (Farley et al. 2018, Nathan et al. 2022, McCrea et al. 2023). High-throughput approaches have enabled the detection of community patterns across broader spatial and temporal scales, while also expanding the taxonomical scope to many previously understudied species groups (Garlapati et al. 2021, van Klink et al. 2022, Abrego et al. 2024).

Nonetheless, novel types of community data are finding their way into experimental studies, too. This development is reflected in the expanding taxonomic scope of experiments conducted during the last decade: bacteria, protists, and fungi are increasingly represented, and these taxa are typically more speciose than macro-organisms such as animals and plants. However, high-throughput technologies still play a rather limited role in experimental community ecology. Some part of this limit to uptake may be due to methodological challenges and costs.

Importantly, issues such as species identification errors and uncertainty (Ficetola et al. 2016), and the inability to quantify abundance reliably (Jo and Yamanaka 2022) may restrict the utility of approaches like eDNA in manipulative studies aiming to test specific hypotheses in community ecology. Nonetheless, the methods for taxonomic assignment are currently developing at a rapid pace (Somervuo et al. 2017, Hleap et al. 2021, Zito et al. 2023, Romeijn et al. 2024, Li et al. 2024) —whereas costs are swiftly decreasing (Ruppert et al. 2019).

While methods are developing, the need for experimental research has remained unchanged. Experiments are still motivated by the goal of testing ecological theory, and the persistent dominance of non-manipulative, observational studies in community ecology adds onus on causal validation. Surprisingly, our review reveals that experiments rarely aim to test hypotheses derived from patterns detected in observational data. This trend may shift as pattern-based knowledge accumulates and high-throughput technologies mature, creating new opportunities for experiments to test and validate the drivers inferred from observed patterns.

The complexity of targeted experimental questions in community ecology has changed over the years. Parallel to an expansion in taxonomic scope (see above), the past decade has seen a shift towards more ambitious experiments in community ecology. An increasing number of experiments now manipulate both biotic and abiotic variables, involve multiple larger taxonomic groups, and/or which score outcomes beyond community structure and function. In particular, we see a growth in the number of experiments focusing on biotic interactions, disease transmission, ecosystem functioning, and evolutionary processes. Although most experiments remain relatively simple, this newly emerging complexity may represent the beginning of a paradigm shift in experimental community ecology. Indeed, such a shift is needed to test predictions emerging from recent advances in theory and inferences derived from the sprawl in observational research. In particular, there is a need for integrating community ecology with other ecological subfields such as ecosystem functioning (Schmitz et al. 2008, Bannar-Martin et al. 2018), host-pathogen interactions (Seabloom et al. 2015), and evolutionary biology (Qian and Jiang 2014, Govaert et al. 2021).

Our review also shows that the community experiments are generally short in duration, as explicitly compared to the generation times of the focal organisms. Where a community experiment will typically span over less than a year, ecological processes often operate over longer timescales. This limitation is especially evident in studies where multiple interacting processes influence the dynamics of interest, but only a subset of these processes is being manipulated or controlled. This challenge can be overcome through long-term experiments (LTEs), which are maintained over decades. Given their longer duration, they are uniquely suited to capture the effects of slow ecological processes and long-term environmental variation. The first experiment of this kind, the Park Grass experiment, was established in 1856 in the UK. While it was originally designed to investigate agricultural productivity, the Park Grass experiment has since become a cornerstone in ecological research. Now spanning over more than one and a half centuries, it has contributed to over 170 scientific publications, and yielded key insights into, e.g., the relationship between abiotic factors, plant richness, and biomass (Silvertown et al. 2006). In the 1980s and 90s, more LTEs have been established, with some of the most well-known being the Cedar Creek LTE (MN, USA), and the Jena experiment (Germany), both aimed at exploring biodiversity-productivity relationships in grasslands. In the Jülich LTE, similar efforts have been made to study the long-term consequences of the order in which species colonize a community for ecosystem functioning (Weidlich et al. 2017). Although there are many more examples of LTEs, they still contribute to the system/taxonomical bias observed in short-term studies. Most LTEs center on grassland communities, with few examples of other systems. Most notably, aquatic ecosystems are underrepresented in LTEs.

Just as temporal scale can constrain inference in experimental research, spatial scale is equally critical. Our review shows that experiments in community ecology are typically done at the local scale, whereas global experiments are rare in comparison. Studying community dynamics at small scales may overlook key context-dependent interactions of biotic processes like dispersal, trophic interactions, and disturbance, which only emerge at broader scales (Winfree et al. 2018). Similarly, abiotic conditions are rarely homogeneous in nature. Therefore, large-scale experiments are usually needed to capture effects of spatial heterogeneity and abiotic gradients (although see Arancibia and Morin 2022, Arancibia 2024 for notable exceptions). Increasing the geographical scale of experiments enables researchers to test whether the observed patterns are consistent across different biogeographical contexts, thus strengthening conclusions.

Clearly, spatial expansion comes with higher costs and increasing logistical challenges. Such restrictions are currently being circumvented by the creation of networks of scientists who together conduct standardized experiments in different geographical locations. Notable examples include the Nutrient Network (NutNet) (Stevens et al. 2015), a consortium coordinating research efforts across more than 130 experimental sites to address questions about environment-productivity-diversity relationships in grasslands. The BugNet (www.bug-net.org) aims to quantify plant and ecosystem responses to herbivores and fungal pathogens through standardized herbivore-exclusion experiments across diverse ecosystems, whereas the International Tundra Experiment (ITEX) (Henry and Molau 1997) coordinates experiments targeting the effects of warming on vegetation across the tundra. The BIODEPTH project (Spehn et al. 2005) aims to experimentally explore the effects of the reduction of biodiversity on ecosystem processes across Europe. Such global networks highlight the potential of coordinated experiments in addressing broad-scale ecological questions.

Each of the experiments described above was conducted in a field setting. Indeed, most experiments in community ecology are conducted in the field, while studies implemented in microcosms and mesocosms remain relatively rare. This scarcity will partly echo past controversies about the value of laboratory studies (Carpenter 1996), as sometimes reflected in funding priorities. Indeed, small-scale experiments allow control over unknown variables and make it easy to create replicate treatments. Both advantages enhance statistical power and clarify underlying mechanisms, making small-scale experiments particularly powerful to test theory. However, this control often limits realism, hindering the extrapolation of the results to natural systems. A more balanced integration of small- and large-scale experimental approaches could enhance the understanding of the causal mechanisms shaping community dynamics.

One approach to addressing current limitations is through whole-ecosystem experiments, where the ecosystem itself serves as the experimental unit, capturing more complex dynamics potentially hidden at the micro- or mesoscales. For instance, in aquatic ecosystems, early small-scale experiments suggested that carbon limitation could control eutrophication in lakes (Schindler 1998). However, whole-lake manipulations later demonstrated that algal blooms can still develop under carbon-limited conditions after phosphorus and nitrogen enrichment. These large-scale experiments revealed that earlier findings were artifacts of the experimental design: they were produced by the absence of CO_2_ exchange in the bottles —a process which occurs naturally in lakes but is absent from bottles (Schindler 1998). This example highlights the importance of whole-scale experiments and how results obtained at a larger scale can then serve as benchmarks for smaller-scale manipulations. Nonetheless, large-scale manipulations can also be logistically challenging, unethical, or impossible to replicate.

In terms of the experimental manipulations applied, our review shows that most experiments rely on press perturbations, while only a small minority use pulse perturbations. This bias in the type of disturbances imposed on communities is relevant because it shapes the inferences that can be made about community interactions (Bender et al. 1984). Sustained press disturbances increase the likelihood of detecting indirect effects (such as trophic cascades, competitive release, etc.). In contrast, pulse perturbations, by allowing us to observe the system after its recovery, are better suited to uncover direct effects. Consequently, the reliance of experiments on press perturbations could bias our perception of the relative importance of direct vs. indirect processes in community dynamics.

A key feature that makes experiments reliable for testing causal patterns is replicability. Experiments allow for replication across multiple experimental units, which strengthens the validity of causal inferences. Our review showed that experiments in community ecology are, however, generally local, short-term, and reliant on manual methods of manipulation and data collection. These limitations of current experimental community ecology can hinder the generalizability of results. Some of these constraints can be overcome by the use of meta-analysis, which synthesizes evidence across experiments to identify general patterns. However, the power of meta-analysis depends greatly on the availability of standardized and highly replicated experimental datasets. The current reliance on manual techniques may reduce the scope for replication, which may limit the utility of synthetic approaches. Importantly, we found that automated methods are virtually never used during the experimental setup and maintenance phase of an experiment. Nonetheless, automation may reduce reliance on human labor, increase the scope for replication, and enable more complex manipulations across both biotic and abiotic axes. In this context, our proof of concept demonstrates how integrating automation into experimental research can significantly advance community ecology.

We conclude that experimental approaches in community ecology need a targeted boost, to efficiently complement observational studies and to test topical theory. To expand the scope, relevance, and impact of experimental research in community ecology, we need to expand both in-situ replication and cross-site reproducibility. These long-lasting challenges can be efficiently addressed by incorporating new technologies in both data collection and processing, and by encouraging collaboration among research groups. The technologies needed are clearly in place, and are increasingly used for biomonitoring (Besson et al. 2022) —whereas new modes of collaboration among tens or hundreds of research teams are expanding the scope for distributed experiments.

## Acknowledgments

All authors thank Mira Kajanus, Ossi Nokelainen, Cara Faillace, Rita Grunberg, members of PredCom (University of Jyväskylä), and the Morin lab (Rutgers University) for their thoughtful comments on multiple drafts of this manuscript. OO and TR were funded by the European Union: The European Research Council (ERC) under the European Union’s Horizon 2020 research and innovation programme (grant agreement No 856506: ERC-synergy project LIFEPLAN). In addition, OO was funded by the Research Council of Finland (grant no. 336212 and 345110), TR by the Swedish Research Council *Vetenskapsrådet* (grant 2023-05118), and NA by the Research Council of Finland (grant no. 342374).

## Conflict of Interest Statement

The authors declare no conflicts of interest related to the contents of this manuscript.

## Notes

http://10.5061/dryad.m905qfvf8

### Competing Interest Statement

The authors have declared no competing interest.

