## Supplementary figure S1 for "Experimental community ecology in decline: A call to embrace technology"

**Appendix S1**

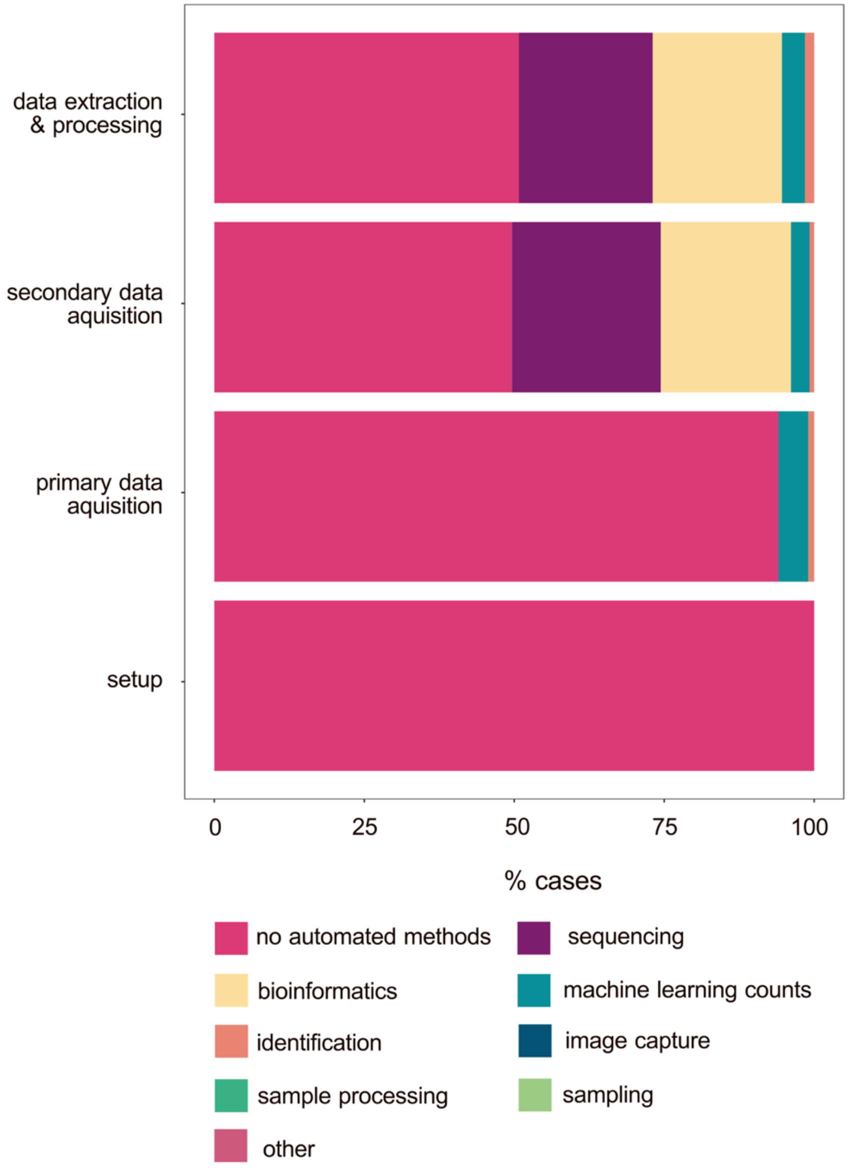


**Figure S1**. Percentage of cases where automated methods were used in experimental community ecology studies 2013-2022.
